# Endogenous pancreatic microRNAs differentially target the Delta, Omicron, and Wuhan SARS-CoV-2 genomes to upregulate the Diabetes-associated genes

**DOI:** 10.1101/2022.01.26.477612

**Authors:** Bhavya Bhavya, Ekta Pathak, Rajeev Mishra

## Abstract

The SARS-CoV-2 viral genome is mutating and evolving into new variations, including the recently discovered Delta and Omicron. In this study, we used and evaluated our notion of differential targeting of SARS-CoV-2 variant genomes by microRNAs (miRNAs) of the infected human pancreas. We found that even with UTR mutations, the Delta, Omicron, and original Wuhan variations’ genomes would be differentially targeted by the host pancreas cell’s native miRNAs in the same way. The miRNAs show a difference in Minimal Free Energy (MFE) with the different viral variants’ genomes; however, they would still be responsible for the upregulation of the diabetes-associated genes.

## Introduction

The SARS-CoV-2 virus is constantly mutating, causing new waves of the COVID-19 pandemic around the world. Until present, the World Health Organization has identified five SARS-CoV-2 virus variants as variations of concern (VOCs), namely Alpha (B.1.1.7), Beta (B.1.351), Gamma (P.1), Delta (B.1.617.2), and Omicron (B.1.1.529)[1]. As of the 21st of January 2022, there had been 340,543,962 confirmed cases of COVID-19 registered worldwide, resulting in 5,570,613 deaths. [2]. In 2022, the most infectious and dominant SARS-CoV-2 variants are the Delta and Omicron being designated as VOCs on 11^th^ May 2021 and 26^th^ November 2021, respectively [1]. The number of mutations in the SARS-CoV-2 genome increased in the Delta and Omicron variants, resulting in lower efficacy of current vaccines based on earlier variations. [3].

In an earlier study, we linked the COVID-19 infection and Diabetes mellitus through differential targeting of the microRNAs (miRNAs) in the human pancreas tissue [4]. We found five Diabetes-associated genes, i.e., CP, SOCS3, AGT, PSMB8 and CFB, to be upregulated in the SARS-CoV-2-infected human pancreas tissue. In the study, we proposed that due to SARS-CoV-2 infection in the human pancreas cell, four host cell’s native miRNAs, i.e., hsa-miR-298, hsa-miR-3925-5p, hsa-miR-4691-3p and hsa-miR-5196-5p, prefer targeting the Untranslated regions (UTRs) of the SARS-CoV-2 genome instead of targeting and regulating the expression of the Diabetes-associated genes. The favorable targeting of the viral genome by miRNAs was supported by the Minimal Free Energy (MFE) that was lower for the hybridization of miRNAs with the viral genome than that with the Diabetes-associated transcripts [4]. This earlier analysis was based on the SARS-CoV-2 genome sequence discovered in Wuhan, China (NCBI Reference Sequence: NC_045512.2) during the pandemic’s initial stages [5]. Here, we sought to see if the earlier hypothesis of differential targeting by miRNAs held true for the genomes of the virus’s Delta and Omicron forms.

## Methods

### Genome UTR sequences of Delta and Omicron variants of SARS-CoV-2

In this study, we obtained the UTR sequences of the Delta and Omicron variant genomes by replacing the mutated nucleotides (mentioned by K. Sato et al. [6]) in the UTR sequences of original Wuhan SARS-CoV-2 genome (NCBI Reference Sequence: NC_045512.2) [5].

### MFE analysis of miRNAs targeting the UTR sequences of SARS-CoV-2 variants’ genomes

The miRNAs targeting the UTR sequences of the Delta and Omicron strain genomes were obtained by using the miRDB online tool [7, 8]. Further, to calculate the MFE of hybridization of miRNAs with the Wuhan, Delta and Omicron genomes, RNAhybrid online tool was used [9, 10]. The MFEs of miRNAs with the UTRs of three viral variant genomes were compared with their already-calculated MFEs with target dysregulated Diabetes-associated transcripts [4].

## Results and Discussion

The Delta variant genome has been shown to be altered at positions 210 (G→T) and 241 (C→T) of the 5’UTR and 29742 (G→T) of the 3’UTR when compared to the UTRs of the original Wuhan variant of SARS-CoV-2. In comparison to the original Wuhan variant UTRs, the Omicron variant genome is only mutated 241 (C→T) position of the 5’UTR[6]. We retrieved the UTR sequences of the Delta and Omicron variant genomes by replacing the aforementioned mutations (Figure 1).

**Figure 1.**
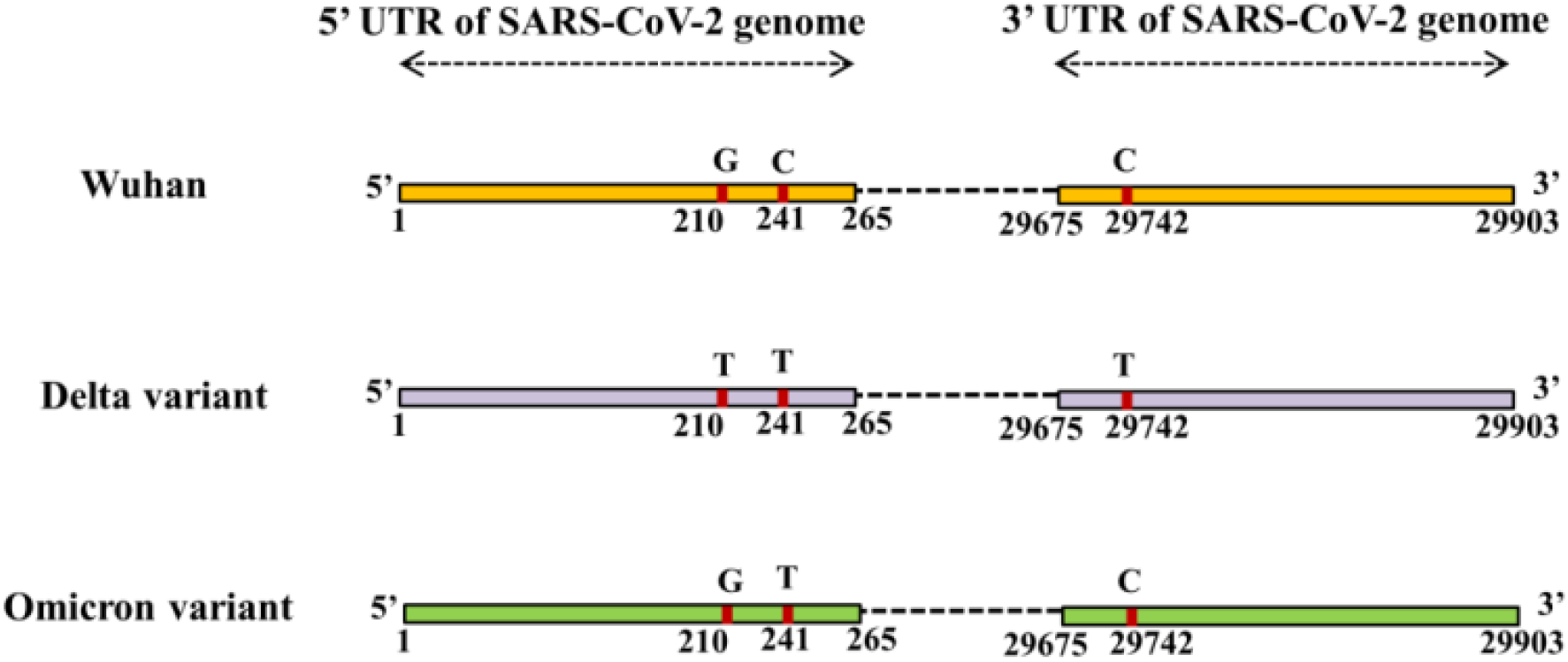
Mutations in the UTRs of the Delta and Omicron variants of the SARS-CoV-2 genome compared to the original Wuhan variant.

The miRNAs targeting the UTR regions of the Delta and Omicron strain genomes were identified using the miRDB online programme [7, 8]. The lists of miRNAs targeting the UTRs of the three virus types were discovered to be identical. In addition, the RNAhybrid online tool was utilised to compute the MFE of miRNA hybridization with the Wuhan, Delta, and Omicron genomes [9, 10]. The calculated MFEs of miRNAs with UTRs from three SARS-CoV-2 variant genomes were compared to the previously calculated MFEs of miRNAs with upregulated Diabetes-associated transcripts (Table 1). Five miRNAs, hsa-miR-8075, hsa-miR-4691-3p, hsa-miR-1283, hsa-miR-4717-3p and hsa-miR-6749-3p, were found to have different MFE with UTRs of different SARS-CoV-2 variants. Among the miRNAs targeting the 5’UTR of the SARS-CoV-2 genomes, hsa-miR-298, hsa-miR-7851-3p, hsa-miR-1303, hsa-miR-3925-5p, hsa-miR-3123, hsa-miR-5196-5p, hsa-miR-4747-5p and hsa-miR-4645-3p show no difference in MFE with the different viral variants. Similarly, hsa-miR-3941, hsa-miR-466, hsa-miR-4775, hsa-miR-5088-5p, hsa-miR-603, hsa-miR-1236-3p, hsa-miR-4279 and hsa-miR-4672 target the 3’UTR of the SARS-CoV-2 genomes and show no difference in MFE with the different viral variants.

**Table 1.** Minimum Free Energy (MFE) of the host pancreas cell’s miRNAs with the SARS-CoV-2 UTRs and dysregulated Diabetes-associated host cell’s transcripts

| miRNAs targeting the 5'UTR of the SARS-CoV-2 genome | Original Wuhan variant genome's 5'UTR | Delta variant genome's 5'UTR | Omicron variant genome's 5'UTR | Dysregulated Diabetes-associated genes in SARS-CoV-2-infected pancreas<br>(Table cells with a dash indicate that the gene transcript is not the target of the corresponding miRNA) |  |  |  |  |
| --- | --- | --- | --- | --- | --- | --- | --- | --- |
|  |  |  |  | CP | SOCS3 | AGT | PSMB8 | CFB |
| hsa-miR-298<sup>#</sup> | -24.8 | -24.8 | -24.8 | - | - | -28.3 | -24.7 | - |
| hsa-miR-7851-3p | -26.7 | -26.7 | -26.7 | - | -30.7 | -29.0 | - | - |
| hsa-miR-1303 | -21 | -21 | -21 | - | -29.9 | -25.5 | - | - |
| hsa-miR-3925-5p<sup>#</sup> | -24.2 | -24.2 | -24.2 | - | -22.9 | - | -22.4 | - |
| hsa-miR-8075* | -25.3 | -25.8 | -25.3 | - | - | - | - | - |
| hsa-miR-4691-3p<sup>#</sup> | -25.5 | -26.2 | -26.2 | -20.7 | -32.1 | - | - | - |
| hsa-miR-1283* | -16.1 | -15.5 | -16.1 | - | - | - | - | - |
| hsa-miR-3123 | -14.7 | -14.7 | -14.7 | -15.8 | -17.0 | - | -15.4 | - |
| hsa-miR-5196-5p<sup>#</sup> | -29.6 | -29.6 | -29.6 | -25.6 | - | - | - | - |
| hsa-miR-4747-5p | -21.3 | 21.3 | -21.3 | -21.3 | - | - | - | - |
| hsa-miR-4645-3p | -20.9 | -20.9 | -20.9 | - | - | -29.0 | - | - |
| miRNAs targeting the 3'UTR of the SARS-CoV-2 genome | Original Wuhan variant genome's 3'UTR | Delta variant genome's 3'UTR | Omicron variant genome's 3'UTR | CP | SOCS3 | AGT | PSMB8 | CFB |
| hsa-miR-3941 | -19.2 | -19.2 | -19.2 | - | -24.1 | - | - | - |
| hsa-miR-466 | -14.7 | -14.7 | -14.7 | -16.2 | - | - | - | - |
| hsa-miR-4775 | -15.7 | -15.7 | -15.7 | -17.3 | - | -19.0 | - | -18.7 |
| hsa-miR-4717-3p* | -26.3 | -27.6 | -26.3 | - | -32.0 | - | - | - |
| hsa-miR-5088-5p | -21.8 | -21.8 | -21.8 | - | -34.6 | - | -26.0 | - |
| hsa-miR-603 | -18.1 | -18.1 | -18.1 | - | - | - | - | - |
| hsa-miR-6749-3p* | -21.4 | -20.5 | -21.4 | - | -35.6 | - | - | - |
| hsa-miR-1236-3p | -24.5 | -24.5 | -24.5 | - | -34.7 | - | -25.7 | - |
| hsa-miR-4279 | -19.3 | -19.3 | -19.3 | - | -31.8 | - | -23.1 | - |
| hsa-miR-4672 | -22.9 | -22.9 | -22.9 | - | - | - | - | - |

\*miRNAs showing difference in MFE with the UTRs of different SARS-CoV-2 variants' genomes.
<sup>#</sup>miRNAs showing lower MFE with the SARS-CoV-2 UTRs than with the transcripts of their target dysregulated Diabetes-associated genes.

hsa-miR-8075 has a lower MFE with the Delta variant’s 5’UTR than with the Wuhan and Omicron variants, which have the same MFE value. hsa-miR-4691-3p had a lower MFE value with the 5’UTR of Delta and Omicron variants than with the Wuhan variant. hsa-miR-1283 has a larger MFE with the Delta variant’s 5’UTR than with the Wuhan and Omicron variants, which have the same MFE value. Because there is no mutation in the 3’UTR of the Omicron variant genome compared to the Wuhan genome, the MFE values for both variants are the same. hsa-miR-4717-3p and hsa-miR-6749-3p have lower and higher MFEs with the Delta variant’s 3’UTR than the Wuhan and Omicron variants.

Among the miRNAs showing a difference in MFE with different variants, only hsa-miR-4691-3p has the MFE having higher value with a Diabetes-associated upregulated gene (i.e., CP) than with the UTR region of any of the three viral variants’ genomes. It shows that even if this miRNA has different MFE with Delta and Omicron variants, then also it favors their genomes over targeting the CP transcript.

Conclusively, we can say that even with the mutations in the genome UTRs, the Delta and Omicron variants of SARS-CoV-2 should lead to a similar phenomenon of differential targeting by miRNAs as mentioned in our earlier study for the Wuhan variant. We propose that if the Delta or Omicron variant of SARS-CoV-2 infects the human pancreas, then this differential targeting by four miRNAs, hsa-miR-298, hsa-miR-3 925-5p, hsa-miR-4691-3p, and hsa-miR-5196-5p, would still be responsible for the upregulation of the diabetes-associated genes, i.e., CP, SOCS3, AGT, PSMB8 and CFB, as was the case with the original Wuhan variant.

## Author contributions statement

R.M conceived and supervised the study. B. performed literature survey, bioinformatic analysis, and prepared the illustrations; B. and R.M wrote the manuscript. E.P contributed in the analysis and manuscript editing. All the authors approved the final version of the manuscript before submission.

## Acknowledgments

Bhavya is supported with a Junior Research Fellowship from the Indian Council of Medical Research (*ICMR*), Government of India, and New Delhi. Research facility support to R.M from the IoE, Banaras Hindu University is also gratefully acknowledged.

